# Competitive approach of invasive cocklebur (*Xanthium strumarium*) with native weed species diversity in Northeast China

**DOI:** 10.1101/2020.01.17.910208

**Authors:** M. F. Iqbal, M. C. Liu, A. Iram, Y. L. Feng

## Abstract

*Xanthium strumarium* is yearly weed local to North America and is presently an obtrusive species. The intrusive weed can contend with local decent variety may turn into a hazardous weed for the agrarian profitability and rangeland biological systems. The present examination researched the challenge of intrusive and local weed populaces inside two straight out variables in matched quadratic rings (treatment/invasive with control/non-invasive). The present investigation was led at four unique areas of northeast China to discover competition of *Xanthium strumarium* on 40 paired matched vegetations in same condition and living space conditions. The number of species (NOS) occurred in quadratic ring; abundance (A); Simpsons Diversity Index (SDI); Margalef’s Richness Index (MRI) compared between invasive and non-invasive quadrates by t-test was recorded significant (P<0.05) suggestion of competitions between plant communities. The abundance in communities decreased significantly in invasive compared to non-invasive quadrate gave an indication about low productivity of plant species due to *Xanthium strumarium*. Rarefaction bend with respect to coefficient of determination (R^2^) explored in the overviewed network (0.86) proposed that there is a solid positive polynomial connection between various weed families. Greatest difference list (87.06%) recorded in Huailai province followed by Yangyuan (44.43%), Zhangjiakou (40.13%) and at Fushun (29.02%). Significant (P<0.05) maximum global R demonstrated high species decent variety was found in Huailai area (0.943) trailed by Zhangjiakou. Significant (P<0.05) density of native weed was recorded in non-invasive quadrate which was comparable to the invasive quadrate. Finally invasive *Xanthium strumarium* compete with native weeds diversity created significant threat to the natural diversity. Most extreme thickness of weed species gave cautioning that the predominant edaphic and natural states of the uneven regions are profoundly favorable for the dispersion and development of the weed in future.

## Introduction

Invasive weeds are presenting critical danger to the biodiversity and biological system working [1–3]. Biological invasions assumed a crucial job in all biological systems [4] however, *X. strumarium* is creating hindrance in the natural vegetations by its competition for food, shelter and light [5]. Biological invasions are considered as succeeding possible hazard to the natural diversity [6]. The invasive weeds reckoned one of the most terrible invaders in the planet characterized to their massive biomass [7,8]. The loss in biodiversity plays a vital role in global climatic change scenario [9] through significant effects on ecosystem [10,11]. These effects adapted drastically by biotic relationship between trophic levels [12,13].

Cocklebur (*Xanthium strumarium* L.) belongs to family Asteraceae are monoecious, annual herbaceous broadleaved, tap rooted, ridged, rough and hairy plant (Venodha, 2016 [14,15] native to North America and Argentina [16–18]. *Xanthium* germinates at broad spectrum temperatures and can invade great areas of marshland and drought resistant species [19]. This weed compete with native weed populations in China are known invader and high potential for colonizing in new areas [20]. *Xanthium* is a harmful invasive weed introduced in Beijing [21] and spread drastically into six provinces [22,23] at China. These provinces are located on the coastal and border area which is previously described the hypothesis that this invasive weed was transported to China by means of intercontinental traffic [22].

This weed basically present in the northeastern territories (Liaoning, Hebei, Beijing and Shandong) having calm mainland overwhelming precipitation with common climate, the southeastern area (Guangdong and Guangxi) is subtropical and northwestern Territory (Xinjiang) having unmistakable mild semiarid particular climatic conditions. Because of wide range climatic conditions, this weed has the ability to adjust essentially to assorted natural condition. Natural displaying has anticipated that this invasive weed have propensity to spread different zones in China [23].

It is a troublesome and problematic noxious weed in many agricultural crops such as maize; common bean; mustard, teff; bread wheat; sweet potato; peanut; barley; chickpea; lentil; Soybean and cotton in many part of the world [5,24]. However the infestation of *X. strumarium* causing the utmost reduction in yield [25]. *X. strumarium* yield losses 31% upto 39% in groundnut crop having four plants in one quadratic ring however 27% yields losses was recorded in maize density 4.7m^−2^ in row planted crop [25]. It invades economic impact on fallow lands resulting in decline the fodder production ultimately affecting in livestock production especially on pig, cattle and sheep [26–28]. The dispersal mechanism of *X. strumarium* was investigated by seed and flood, animals, wind, vehicles and human clothing however its long dormancy period may be helpful for its pressure [5].

Native plants are important in rangeland ecosystems that evolved to grow in local conditions, however then even do not require any specific irrigations and foliar nutrients. The leaves of the native plants have high amounts of inexpensive source of nutrients for human as well as animal consumptions in developing countries [29].

The aim of this study is to identify invasive and native weeds and to predict the appropriate habitation and ecological distribution of *X. strumarium.*

The research highlights and hypothesis are as under:

1. Impact of invasive cocklebur in native diversity of weed in natural ecosystem
2. Maximum weed density gave the indication that prevailing environmental conditions of the vicinity are conducive for the distributions of invasive weeds.
3. The impact of *Xanthium strumarium* invasion was investigated at different locations.
4. Co-efficient of determination (*R*^2^) suggested strong positive relationship between different families in ecosystem.

The field survey conducted to evaluate the impact and competitions of invasive and non-invasive weeds present in selected quadratic rings at different sites in same habitat and environmental cues under Liaoning Key Laboratory for Biological Invasion and Global Changes, College of Bioscience and Biotechnology, Shenyang Agricultural University, Shenyang, Northeast China.

## Materials and Methods

The present study was conducted during peak months (August-September) in four different locations of northeast China by line transects ecological method [30] during 2018 to investigate competitions of *Xanthium strumarium* and native weeds present in invasive and non invasive quadrate (Figure 1). These locations (Fushun County, Zhangjiakou, Huailai County, and Yangyuan County), however, Fushun County located in Liaoning Province with comparatively high precipitation (mm); however, the other three locations situated in Hebei Province located between Beijing and Inner Mangolia having comparatively 50% low precipitation (Figure 2). Due to dynamic climate conditions and combination of hilly areas, these regions are rich biodiversity of native and invasive populations of weeds.

**Figure 1.**
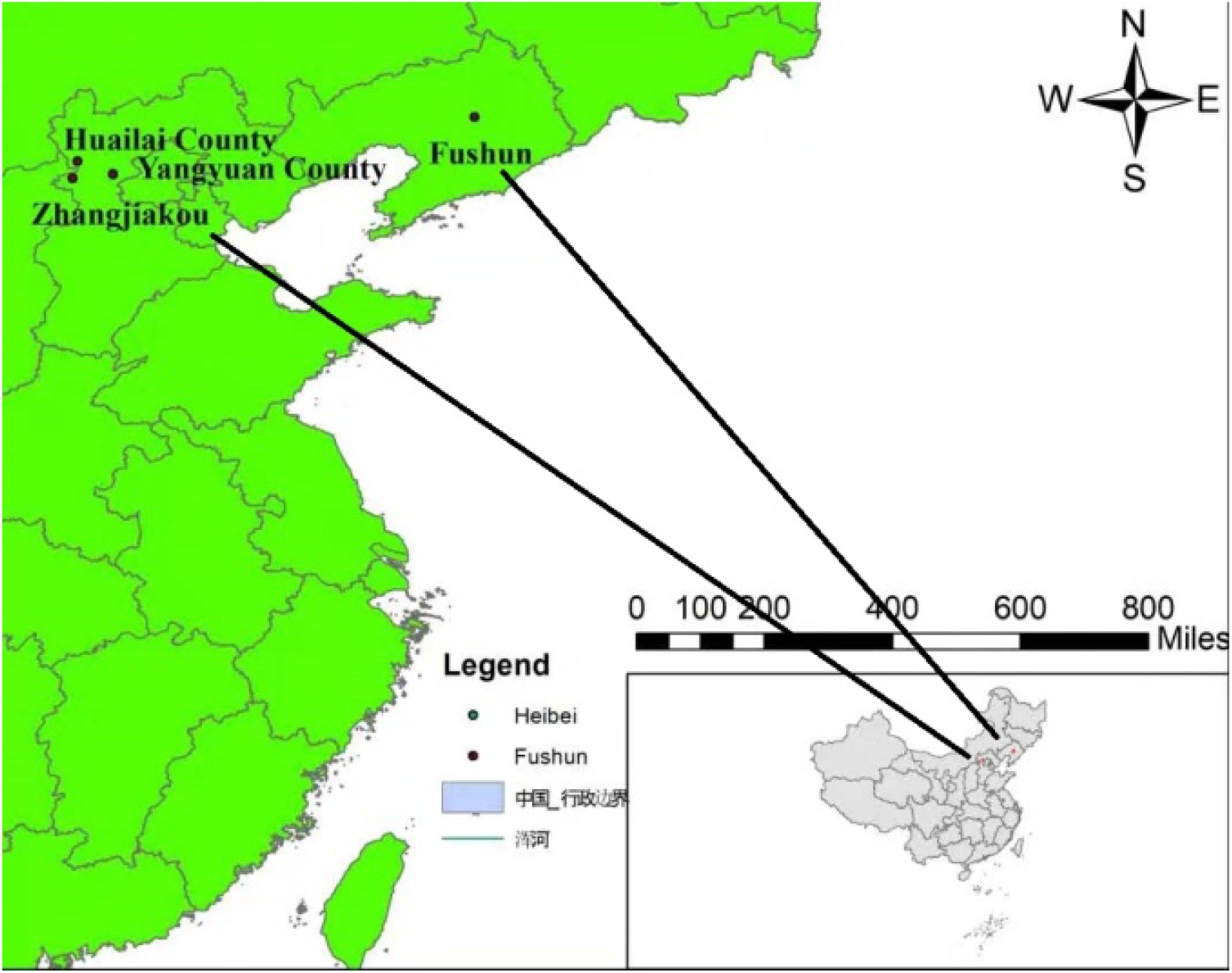
Latitudinal map showing different study sites in Northeast China in 2018 [32,57]

**Figure 2.**
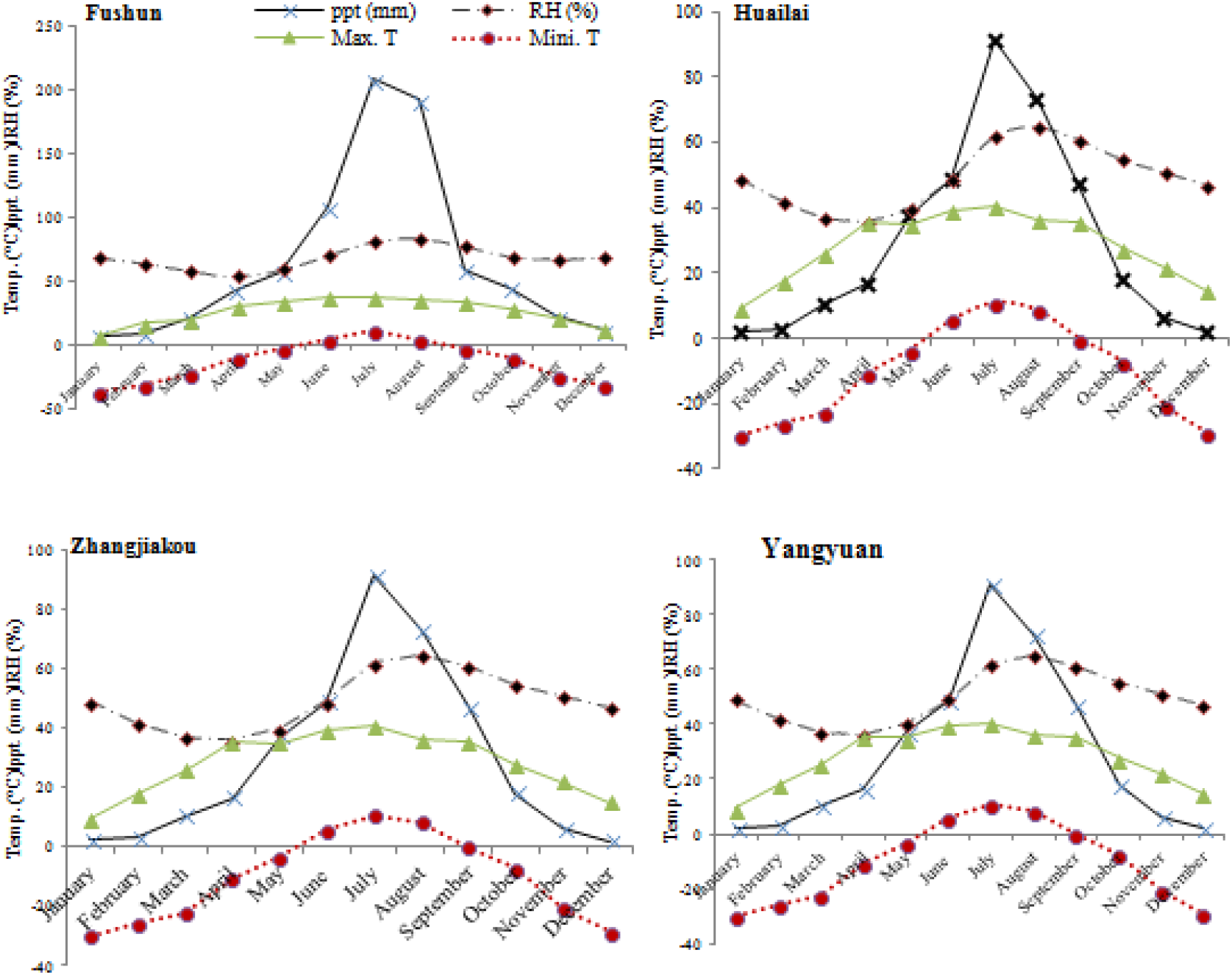
Monthly climatic data of different locations at Liaoning and Hebei Province for the year 2018

The two factors were studied (Invasive versus nearby site non-invasive) keeping in view identical ecological circumstances having 20m in a radius [31]. On visual observations where the *Xanthium strumarium* showed its dominance considered as treatment (Invasive) and 2^nd^ vegetation plot selected near invasive treatment considered as the non-invasive/control [19]. Ten pairs of plots selected from each site and sum 40-paired plots investigated. In each selected site, ten quadrates each having 100×100cm size selected and all the individual weed populations recorded in both invasive and non-invasive quadrates (Table 1). This procedure was repeated three times in a site to reduce confounding factors. The geographic coordinates of four different paired sites (Figure 1) were recorded by GPS system and ArcGIS version 9.2 software (ESRI) used to map out the distribution of studied sites [32].

**Table 1.** Plant species found during extensive field survey with family, habitat, and life form

| **Sr. No.** | **Species** | **Local Name** | **Families** | **Abb.** | **Habitat Range** | **Life form** |
| --- | --- | --- | --- | --- | --- | --- |
| 1 | *Xanthium strumarium* | Common cocklebur | *Asteraceae* | *Xs* | *Invasive* | *Herb* |
| 2 | *Humulus scandens* | Wild hop | *Cannabaceae* | *Hs* | *Native* | *Herb* |
| 3 | *Setaria feberi* | Chinese foxtail | *Poaceae* | *Sf* | *Native* | *Grass* |
| 4 | *Chenopodium album* | Bathu | *Amaranthaceae* | *Ca* | *Native* | *Herb* |
| 5 | *Artemisia annua* | Annual wormwood | *Asteraceae* | *Ara* | *Native* | *Herb* |
| 6 | *Dioscorea zingiberensis* | Yellow ginger | *Dioscoreaceae* | *Dz* | *Native* | *Herb* |
| 7 | *Setaria pumila* | Yellow foxtail | *Poaceae* | *Sp* | *Native* | *Grass* |
| 8 | *Erodium stephianum* | Pin clover | *Geraniaceae* | *Es* | *Native* | *Herb* |
| 9 | *Seteria viridis* | Green foxtail | *Poaceae* | *Sv* | *Native* | *Grass* |
| 10 | *Malva verticillata* | Chinese mallow | *Malvaceae* | *Mv* | *Native* | *Herb* |
| 11 | *Avena sativa* | Common oat | *Poaceae* | *As* | *Native* | *Grass* |
| 12 | *Cynanchum chinensis* | Dog strangling vine | *Apocynaceae* | *Cc* | *Native* | *Herb* |
| 13 | *Artemisia selengensis* | Wormwood | *Asteraceae* | *Ars* | *Native* | *Herb* |
| 14 | *Oxalis corniculata* | creeping woodsorrel | *Oxalidaceae* | *Oc* | *Native* | *Shrub* |
| 15 | *Cynanchum wilfordii* | Baekshuoh | *Apocynaceae* | *Cw* | *Native* | *Herb* |
| 16 | *Acalypha australis* | Asian copperleaf | *Euphorbiaceae* | *Aca* | *Native* | *Shrub* |
| 17 | *Taraxacum mongolicum* | Dandelion | *Asteraceae* | *Tm* | *Native* | *Herb* |
| 18 | *Myosoton aquaticum* | Giant chickweed | *Caryophyllaceae* | *Ma* | *Native* | *Herb* |
| 19 | *Polygonum aviculare* | Common knotweed | *Polygonaceae* | *Pa* | *Native* | *Herb* |
| 20 | *Ambrosia trifida* | Giant ragweed | *Asteraceae* | *Amt* | *Invasive* | *Herb* |
| 21 | *Ambrosia artemisifolia* | Common ragweed | *Asteraceae* | *Ama* | *Invasive* | *Herb* |
| 22 | *Axonopus compressus* | Cow Grass | *Poaceae* | *Axc* | *Native* | *Grass* |
| 23 | *Medicago lupulina* | hop clover | *Fabaceae* | *Mel* | *Native* | *Herb* |
| 24 | *Erigeron acris* | bitter fleabane | *Asteraceae* | *Ea* | *Native* | *Herb* |
| 25 | *Euphorbia maculata* | prostrate spurge | *Euphorbiaceae* | *Eumc* | *Native* | *Shrub* |
| 26 | *Leptochloa chinensis* | Chinese sprangletop | *Poaceae* | *Lc* | *Native* | *Grass* |
| 27 | *Plantago depressa* | broad-leaved plantain | *Plantaginaceae* | *Pd* | *Native* | *Herb* |
| 28 | *Rubia cordifolia* | common madder | *Rubiaceae* | *Rc* | *Native* | *Herb* |
| 29 | *Cyperus cyperoides* | Madhana | *Cyperaceae* | *Cc* | *Native* | *Sedge* |
| 30 | *Kochia scoparia* | Kochia | *Chenopodiaceae* | *Ks* | *Native* | *Herb* |
| 31 | *Chenopodium hybridum* | Maple-leaved goosefoot | *Amaranthaceae* | *Ch* | *Native* | *Herb* |
| 32 | *Chenopodium ficifolium* | Fig-leaved goosefoot | *Amaranthaceae* | *Cf* | *Native* | *Herb* |
| 33 | *Eleusine indica* | yard-grass | *Poaceae* | *Ei* | *Native* | *Grass* |
| 34 | *Polygonum longisetum* | knotweed | *Polygonaceae* | *Pl* | *Native* | *Herb* |
| 35 | *Echinochola crus-galli* | cockspur grass | *Poaceae* | *Ecg* | *Native* | *Grass* |
| 36 | *Zizania latifolia* | Zizania | *Poaceae* | *Zl* | *Native* | *Grass* |
| 37 | *Imperata cylindrica* | kunai grass | *Poaceae* | *Ic* | *Native* | *Grass* |
| 38 | *Amaranthus retroflexus* | rough pigweed | *Amaranthaceae* | *Ar* | *Native* | *Herb* |
| 39 | *Amaranthus blitum* | Guernsey pigweed | *Amaranthaceae* | *Ab* | *Native* | *Herb* |

Bar graph regarding rarefaction described the number of individual species at species level suggested species richness in the studied sites calculated by results of samplings. After collecting the species population data were focused on multivariate analysis with invasive and non invasive quadrate (used factor) using PRIMER 7 software [33], this data sets was used to ordinate the similarity between these quadrates based on Bray-Curtis dissimilarity [31]. The competition was calculated by similarity analysis and percentage [34,35]. According to these techniques the impact of each weed species contributing to dissimilarity between groups were calculated [36]. Margalef’s Richness Index (MRI) was calculated by counting one and sum total of all species in ten quadrates (Sinha, 2017), Simpson Diversity Index (SDI) was also calculated at species level [31,37]. The abundance and density of each weed recorded in studied quadratic rings by dividing it with sum total of quadratic rings studied, however number of species were calculated directly by counting method [31,38–42]. Comparison of number of weed species present in invasive and non invasive quadrate was calculated [43]. The total weed species present in a selected community subjected to ANOVA and the mean difference between Invasive and non-invasive quadrate calculated by t-test at 5% level of significance.

## Results

The rarefaction curve indicated that the individuals of Asteraceae family are more diversified showed highly positive polynomial *Coefficient of determination R*^2^ = 0.88 compared to other species studied (Figure 3). During this survey sum of 15 plant species from 9 genera were found, however 10 species were recorded in non invasive quadrate and 12 from the plots with invasive quadrate with varying number of individuals (Table 1). Significant high (P<0.05) number of species investigated at Fushun (Figure 4A); Zhangjiakou (Figure 4C) and Yangyuan (Figure 4D) locations in non invasive quadratic ring compared to invasive. However, significant (P<0.05) maximum weed species abundance was recorded at Fushun (4.00); Huailai (9.00) and Yangyuan county (3.00) in non invasive quadratic ring. Other factor regarding Simpson’s Diversity Index (SDI); Margalef’s Richness Index (MRI) gave the clear-cut indication that *Xanthium strumarium* present in invasive quadrate compete positively. Maximum SDI investigated higher diversity index or probability between invasive and non invasive quadratic rings, however, 0.99 and 0.97 values indicated more diversified species in the study areas. Significant (P<0.05) result in weed density was recorded in non-invasive quadratic ring. On the other side invasive quadrate investigated non-significant (P<0.05) effect and recorded high competition in density between two groups of weeds at Zhangjiakou (Figure 5A). In Huailai county significant (P<0.05) result was investigated in non-invasive quadratic ring. The density of weed populations in invasive quadratic ring differed non significantly (Figure 5B). In yangyuan, the density of non-invasive quadrate differed significantly (P<0.05) with invasive ring (Figure 5C). Same trend but highly significant density was investigated in non-invasive quadratic ring, which was comparable to the invasive quadrate. Maximum dissimilarity index (87.06%) recorded in Huailai county followed by Yangyuan (44.43%), Zhangjiakou (40.13%) and at Fushun (29.02%). Significant (P<0.05) maximum global R indicated high species diversity found in Huailai county (0.943) followed by Zhangjiakou. Similarity (%) data analysis described that weeds computed average dissimilarity with invasive and non-invasive quadrates (Table 2). SIMPER analysis pooled data based on average abundance of each weed species illustrated that *Malva verticilata* (10.40) contributed maximum in non-invasive quadrate followed by *Chenopodium album* (9.00); Avena sativa (7.10) and *Seteria viridis* (6.80) compared to invasive quadrate at Yangyuan County. However, *Malva verticilata* (23.94%) contributed maximum both quadratic rings compared to *Seteria viridis* (20.80%). *Seteria viridis* (14.00%; 40.82%) contributed maximum at Huailai and Fushun area followed by *Seteria febrei* (24.40%) at Zhangjiakou. A total of 12 weeds species in different families documented from invasive and non invasive forty paired quadratic rings in different studied area. These results clearly indicate that at first level Primer divided the weed populations of the whole study area into one major community at association level and that further divided into two sub communities at sub association level (Table 1). The similarity analysis found significant degree of variation between species compositions in invasive & non invasive quadrates having global statistics R values of 0.943 (P<0.01), 0.779 (P<0.01), 0.62 (P<0.01), 0.553 (P<0.01) for Huailai, Zhangjiakou, Yangyuan and Fushun. Similarity (%) data analysis described that weeds contributing average dissimilarity in invasive and non-invasive quadrates (Table 2). A shade plot showing the multivariate analysis and magnitudes of the invasive and non-invasive weeds distributions. However, the each color represents the species present in an ecosystem. Hierarchical data showed gave an indication of total number of species studied in four different dimentional community (Figure 6).

**Figure 3.**
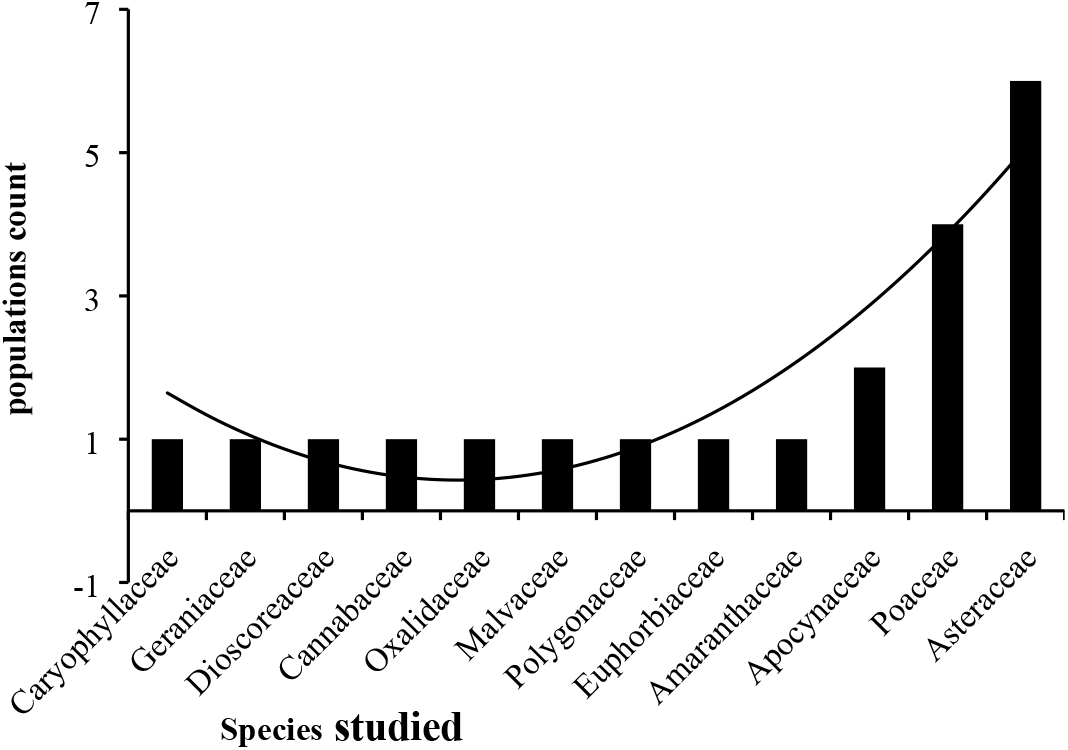
Rarefaction curve showing cumulative number species studied at four locations with *Coefficient of determination* (*R*^2^)

**Figure 4.**
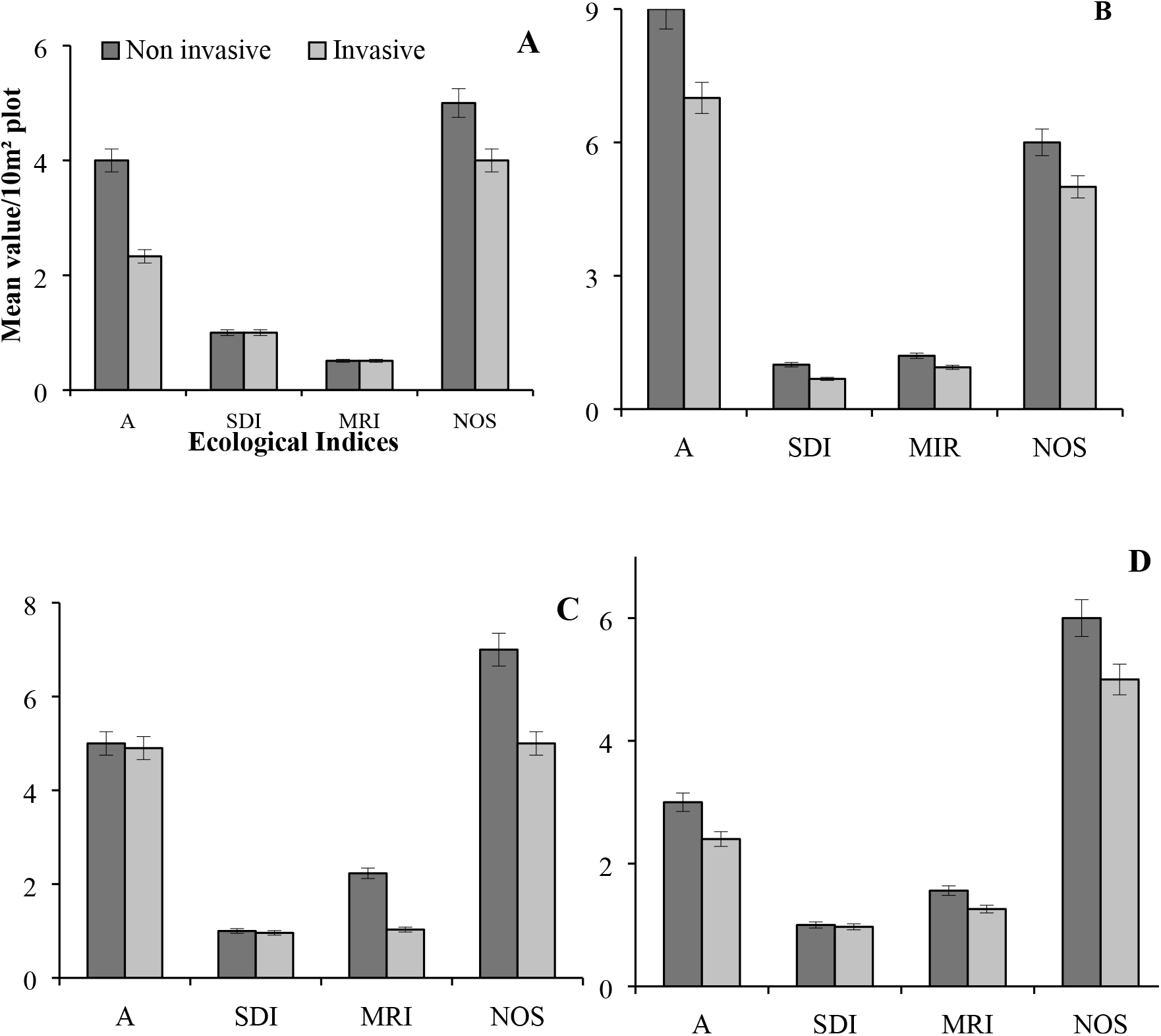
showing ecological indices regarding mean value 10 m^−2^ of Invasive with Non Invasive quadratic rings at different sites where A (Fushun); B (Huailai); C (Zhangjiakou); D (Yungyuan). Whereas (A = Abundance; SDI = Simpsons Diversity Index; MRI = Margalef’s Richness Index; NOS = Number of Species).

**Figure 5.**
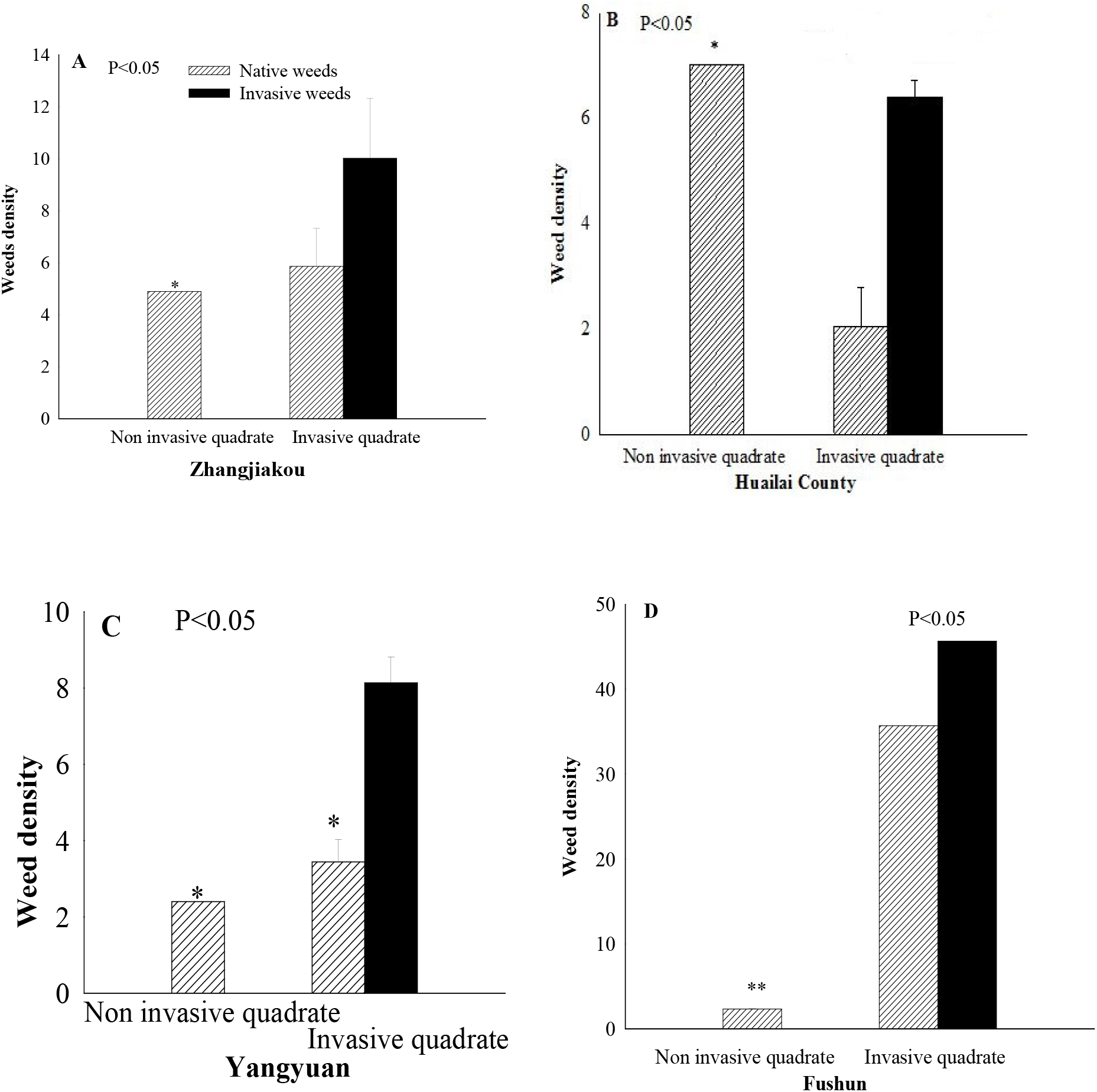
showing weed density 10 m^−2^ of invasive with non invasive/control quadrate at different locations (A) Zhangjiakou; (B) Huailai County; (C) Yangyuan county; (D) Fushun County

**Fig. 6.**
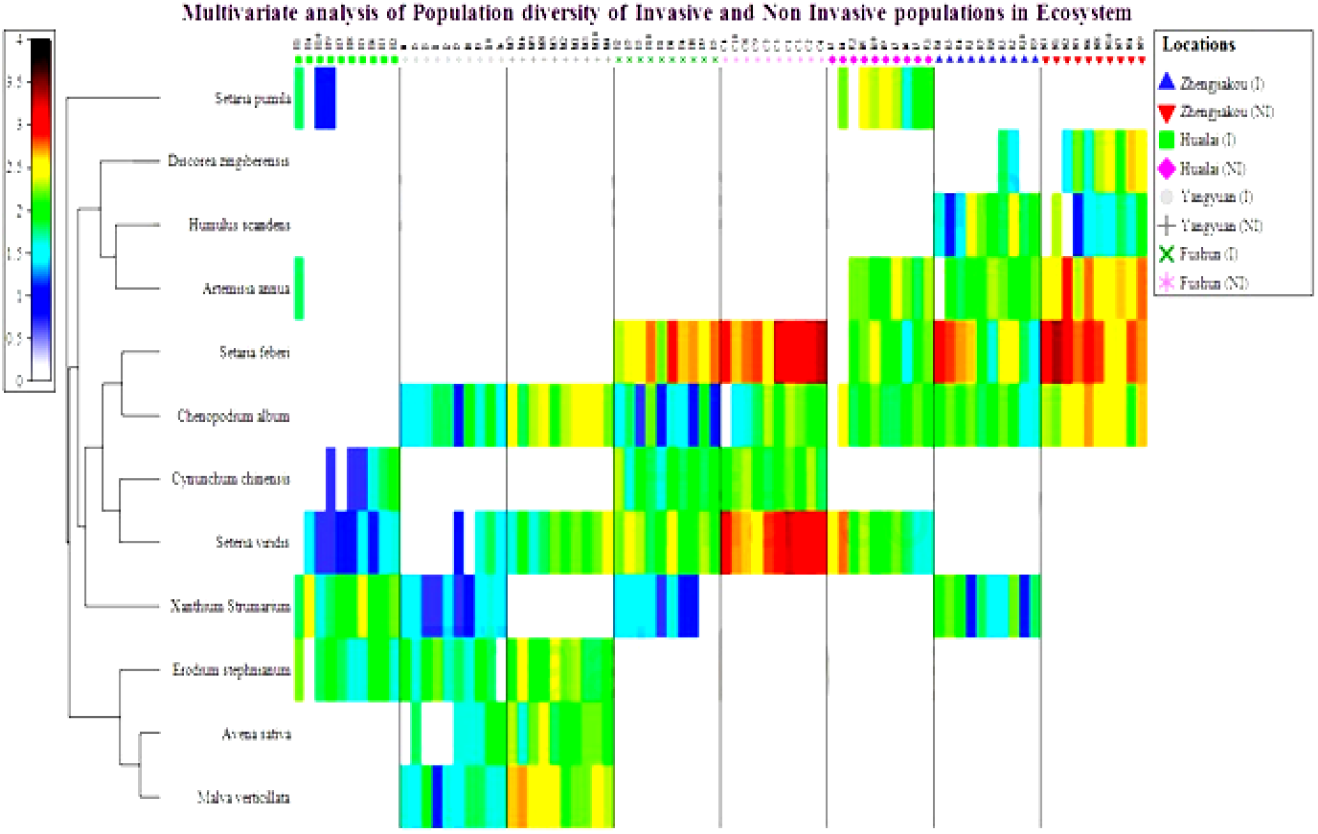
A shade plot showing the multivariate relative magnitudes of the invasive and non-invasive weeds distributions where the color shading represents the population diversity in a community. Hierarchical data showed that total number of species studied during extensive field survey during 2018

**Table 2.** Pooled data of SIMPER analysis of *Xanthium strumarium* invasive and non-invasive/control quadrate in four locations at Northeast China

| Average dissimilarity = 44.43% based on Average abundance (Yangyuan County) |  |  |  |  |  |
| --- | --- | --- | --- | --- | --- |
| Species | Invasive | Non-Invasive | Av. Diss. | Diss./SD | Contribution (%) |
| *Malva verticilata* | 4.00 | 10.40 | 10.64 | 2.54 | 23.94 |
| *Seteria viridis* | 1.40 | 6.80 | 9.24 | 1.95 | 20.80 |
| *Chenopodium album* | 4.10 | 9.00 | 8.27 | 2.27 | 18.62 |
| *Avena sativa* | 2.90 | 7.10 | 7.38 | 1.48 | 16.62 |
| Average dissimilarity = 87.06% based on Average abundance (Huailai County) |  |  |  |  |  |
| *Seteria viridis* | 2.10 | 7.60 | 12.19 | 1.29 | 14.00 |
| *Chenopodium album* | 0.00 | 6.10 | 12.05 | 2.14 | 13.84 |
| *Artemisia annua* | 0.50 | 6.10 | 11.53 | 1.69 | 13.24 |
| *Setaria pumila* | 0.90 | 6.50 | 11.46 | 1.78 | 13.16 |
| Average dissimilarity = 40.13% based on Average abundance (Zhangjiakou) |  |  |  |  |  |
| *Seteria febrei* | 10.00 | 16.80 | 9.79 | 1.30 | 24.40 |
| *Discorea zingiberensis* | 0.80 | 6.50 | 7.28 | 1.44 | 18.13 |
| *Artemisia annua* | 6.30 | 12.50 | 7.27 | 1.81 | 18.12 |
| Average dissimilarity = 29.02% based on Average abundance (Fushun County) |  |  |  |  |  |
| *Seteria viridis* | 7.90 | 17.50 | 11.84 | 2.41 | 40.82 |
| *Seteria febrei* | 12.30 | 18.10 | 7.86 | 1.47 | 27.10 |
| *Chenopodium album* | 3.60 | 5.80 | 4.31 | 1.58 | 14.86 |

## Discussion

The data of study showed that invasive weed (*Xanthium strumarium*) investigated at these locations having maximum competition with native weeds. Significant competition of native weeds recorded in non-invasive quadrate recorded high significant abundance at Huailai County. However, on all other three sites native weeds compete with invasive weed that was comparable to non invasive quadratic ring. Simpson’s Diversity Index (SDI); Margalef’s Richness Index (MRI) gave the clear-cut indication that *Xanthium strumarium* present in invasive quadrate compete positively within non-invasive weeds. Maximum SDI investigated higher diversity index or probability between invasive and non-invasive quadratic rings. These results were in line with the scientists who reported that *Xanthium strumarium* affected severally to the natural vegetations. Seventy species investigated in the non-invasive plots, however, thirty-one recorded in *Xanthium strumarium* toxic area. About 55.71% species reduced in invasive compared to non-invasive range. SDI, MRI reduced upto 70.33%, and 69.39% in the invasive plot [28,31]. The results of our study was also in line with the researchers who studied that the effect of invasive species on native population was usually stronger than vice versa [44]. Comparatively significant species diversity were present in Invasive quadrate in Huailai (4B) followed by Zhenjiakou (4C); Fushun (4A) and Yangyuan (4D) than non invasive quadrat that prove the hypothesis that Invasive species compete with the native species abundance because they have high phenotypic plasticity and have maximum ability to adopt and enhance vegetative growth in an ecosystem.

These results are in accordance to the researchers who reported that *X. strumarium* is an invasive possessing negative impacts on natural ecosystem [45]. This weeds can alter abundances of native weeds in a terrestrial food web, but the pattern is complex [28,46,47]. *Xanthium strumarium* having a big canopy and tap root system caused considerable impact on native plant communities; however, its population abundance causes serious threat to the native plant communities. Maximum evenness recorded in invasive quadrate investigated good competitor in ecosystem processes. The plant invasions severely influence on the size and composition of native vegetations through interference of biotic interactions in abiotic network [48,49]. *X. strumarium* caused maximum damage to the native weeds might continue in near future [5,50]. Invasive species are characterized in wide-ranging, rapacious, assertively persistent, elastic, grow rapidly, capability to travel in extensive areas and rich reproduction as these flora emerge to have particular qualities that permit them to compete with native plant species [51,52]. The coefficient of determination (*R^2^*) recorded in the surveyed community of different weeds families suggested that there is a strong positive polynomial relationship investigated at different locations. These finding are in accordance to the other studies on invasive weeds of Asteraceae family in which the researcher indicated the strong effect of invaders on the ecosystem properties [31,53]. The researchers deliberated that *Xanthium strumarium* became most brutal Invasive Alien plant dominated on rangelands of Ethiopia [54]. The researchers previously conducted paired wise survey in invasive and non invasive quadrate suggested that invasive plant species compete and affect positively on native plant species [55]. Our hypothesis is true that Invasive weeds have a potential to compete with of non-invasive/native weeds species in a community. These results were in line to the researchers who reported heterogeneous population of weed community in their experiments [38]. Maximum dissimilarity index (87.06%) was recorded in Huailai county followed by Yangyuan (44.43%), Zhangjiakou (40.13%) and at Fushun (29.02%). *Xanthium strumarium* affected on the species density in our experimental study which were in line with the researchers who described the Jaccard’s similarity index between invasive and non-invasive ranges in weed communities which gave indication about 38.4% loss of weed species due to *Xanthium strumarium* infested plot resulted 61.6% dissimilarity index [28]. In our experiment significantly (P<0.01) maximum global test for sample statistics R values were recorded in Huailai County (0.943) followed by Zhangjiakou, Yangyuan and Fushun area investigated by analysis of similarities using one way analysis of variance taking 999 number of permutations. The larger value of R indicated that higher effect of species diversity upon studied variables. Huailai County significantly more affected due to *Xanthium strumarium* invasions compared to Zhangjiakou and Yangyuan areas. The lowest invasion was investigated in Fushun because of highest dissimilarity or beta diversity in invasive and non invasive/control plots. The result of our experiment suggested that there is a strong negative relationship (*R^2^* =0.741, or 74.1%) found between the *Xanthium strumarium* (%) and Species Richness. These results are in line with the scientists who reported that regression equation and Pearson correlation (−0.861) indicated the existence of strong negative linear relationship between *Xanthium strumarium* and Richness however with the increase in *Xanthium* species (%) the richness decreases significantly [28]. In our study, maximum density of natural weed was gave indication that the prevailing edaphic and environmental conditions of the area are highly conducive for the distribution and growth of the weed species. These work is in line with the researchers who reported same results in their experiments [53,56].

## Conclusion

It was reasoned that *Xanthium strumarium* rival local weeds species in a network. Noteworthy challenge in thickness recorded in non invasive quadrate that demonstrate the speculation that invasive weed influenced the local weeds decent variety thought about a huge risk to the characteristic biological system. Greatest SDI at Huailai District gave a sign that weed networks were increasingly heterogeneous in non-invasive locales contrasted with invasive ones. The bigger estimation of global R in measurements demonstrated that higher impact of species decent variety upon examined factors. The most influenced site found by *Xanthium strumarium* attack was Huailai Area followed by Zhangjiakou and Yangyuan separately. The most minimal attack impacts were examined in Fushun in view of most elevated difference or beta assorted variety. Most extreme thickness of weed species gave cautioning that the predominant edaphic and ecological states of the territory are exceptionally favorable for the conveyance and development of the weed species. At long last it is an admonition circumstance to the scientists that there is a critical need of appropriate intending to oversee and dispose of the spread of this weed attack. We also encouraged researchers to explore the defensive mechanism, nutrient cycling, role of pseudomonas bacteria, resistance mechanism of invasive and native weeds in future.

## Author Contributions

The following statements stated that “conceptualization, Y.L.F. and M.F.I.; methodology, Y.L.F.; software, M.F.I.; validation, Y.L.F., M.F.I.; formal analysis, M.F.I.; investigation, M.F.I., L. M. C.; resources, Y.L.F.; data curation, M.F.I., A. I.; writing— original draft preparation, M.F.I.; writing—review and editing, M.F.I; Y.L.F.; visualization, M.F.I.; supervision, Y.L.F.; project administration, Y.L.F.; funding acquisition, Y.L.F.”,

